# Spage2vec: Unsupervised detection of spatial gene expression constellations

**DOI:** 10.1101/2020.02.12.945345

**Authors:** Gabriele Partel, Carolina Wählby

**Affiliations:** Centre for Image Analysis, Dept. of Information Technology and SciLifeLab, Uppsala University, Uppsala, Sweden

## Abstract

Investigation of spatial cellular composition of tissue architectures revealed by multiplexed in situ RNA detection often rely on inaccurate cell segmentation or prior biological knowledge from complementary single cell sequencing experiments. Here we present spage2vec, an unsupervised segmentation free approach for decrypting the spatial transcriptomic heterogeneity of complex tissues at subcellular resolution. Spage2vec represents the spatial transcriptomic landscape of tissue samples as a spatial functional network and leverages a powerful machine learning graph representation technique to create a lower dimensional representation of local spatial gene expression. We apply spage2vec to mouse brain data from three different in situ transcriptomic assays, showing that learned representations encode meaningful biological spatial information of re-occuring gene constellations involved in cellular and subcellular processes.

## INTRODUCTION

Recent advances in single-cell RNA (scRNA) sequencing [1,2] allow to dissect the cell type heterogeneity of complex tissues at incredible pace. An international effort has started building comprehensive reference maps of gene expression at cellular resolution to uncover the cell type composition of entire organs and organisms [3]. However, in order to understand the functional architecture of a tissue it is essential to reconstruct the spatial organization of its constituent cell types. To this end, single cell sequencing analyses are often complemented with imaging-based methods for spatially resolved multiplexed in situ RNA detection [4–8] that allow to map mRNA molecules directly in tissue samples and identify specific cell type location, enabling the discovery of their functional role inside the tissue architecture.

Previous attempts to map the spatial heterogeneity of cell types mostly relied on cell body segmentation algorithms and gene assignments to cells based on segmented cell boundaries [4–7]. Extracted per-cell gene expression profiles are successively clustered and annotated based on complementary scRNA sequencing analysis experiments or published literature [4–7]. This means that analysis of the spatial heterogeneity in tissue samples is limited by the accuracy of image segmentation algorithms to outline exact cell borders in dense and overlapping cell environments, with uneven illumination conditions and low-signal to noise ratios. Moreover, while some cell types are defined by clear differences in their gene expression profiles, others differ by only a few genes in their transcriptome (e.g. like finely related neuronal subtypes) making their identification challenging.

Preliminary work from *Park J, Choi W. et al.* [9] tries to address these problems proposing a segmentation-free spatial cell-type analysis (SSAM) based on cellular mRNA density estimation via Gaussian KDE [10], defining cell location as local maxima of mRNA-dense regions and extracting gene expression profiles for each cell (i.e. local maxima) as the averaged gene expression in that unit area. *Qian X. et al*. [11], instead, proposed a probabilistic framework for jointly assigning mRNAs to segmented cells and cells to cell types based on scRNA-seq cell-type priors, achieving a fine classification of interneurons subtypes of CA1 hippocampal region.

Despite these efforts for improving cell type identification in situ, spatial cell type analyses alone do not use the full power of in situ spatial transcriptomics: The subcellular resolution can reveal spatial heterogeneity also at subcellular levels. There is compelling evidence that many genes are expressed in a spatially dependent fashion independent of cell types [12], and this information is lost when analysing transcriptional profiles of single cells. Moreover, there is a considerable amount of heterogeneity within each cell type explained by the balance between intrinsic regulatory networks and extrinsic subcellular processes depending on the local cellular microenvironment [13–17]. mRNA localization plays an important role in these cell differentiation processes as localization can vary during specific stages of cell development, and distinguishes cell phenotypes, activities and communication. Specifically, mRNA localization is involved in cellular compartmentalization of gene expression into spatial functional domains involved in spatially targeted segregation of protein synthesis [18]. For example, mRNA localization is particularly diffused in neurons, where protein synthesis can take place at distal sites far away from the nucleus: Dendritic and axonal structures express several forms of plasticity that requires local translation [19–22]. Disruption of these subcellular biological processes were shown to be implicated in neurodevelopmental, psychiatric or degenerative diseases [23–26]. It is thus important to take advantage of in situ mRNA detection methods to dissect the spatial heterogeneity of gene expression at subcellular resolution with respect to development and disease, and unreveal the subcellular spatial domains underlying cell differentiation.

Here we propose a novel segmentation free approach for analyzing the spatial heterogeneity in gene expression of tissue samples that does not rely on the definition of cell types and cell segmentation but leverages the spatial organization of single mRNAs to define subcellular spatial domains involved in cellular differentiation. Specifically, we consider the spatial organization of mRNAs inside tissues as a spatial functional network where different mRNA types interact based on their spatial proximity [Figure 1], and where subcellular domains can be identified as clusters of local gene constellations that are shared or cell-type specific. In order to investigate the spatial mRNA network for recurrent gene constellations, we adopted a powerful graph representation learning technique [27] based on graph neural networks (GNN) [28], that has recently emerged as state-of-the-art machine learning technique for leveraging information from graph local neighborhoods. Therefore, each mRNA location is encoded in a graph as a node with a single feature representing the gene it belongs to and it is connected to all the other nodes representing the other mRNAs located in its neighborhood [Figure 1a]. During training, the GNN learns the topological structure of each node’s local neighborhood as well as the distribution of node features in the neighborhood (i.e. local gene expression), and projects each node in a lower dimensional embedding space that encapsulates high-dimensional information about the node’ s neighborhood [Figure 1b]. We call this vectorization approach spatial gene expression to vector, or spage2vec, where geometric relations in this lower dimensional space corresponds to higher order relationships in the local gene environment. We apply spage2vec to three publicly available datasets and compare the resulting gene constellations to cell type maps presented in the respective publications.

**Figure 1.**
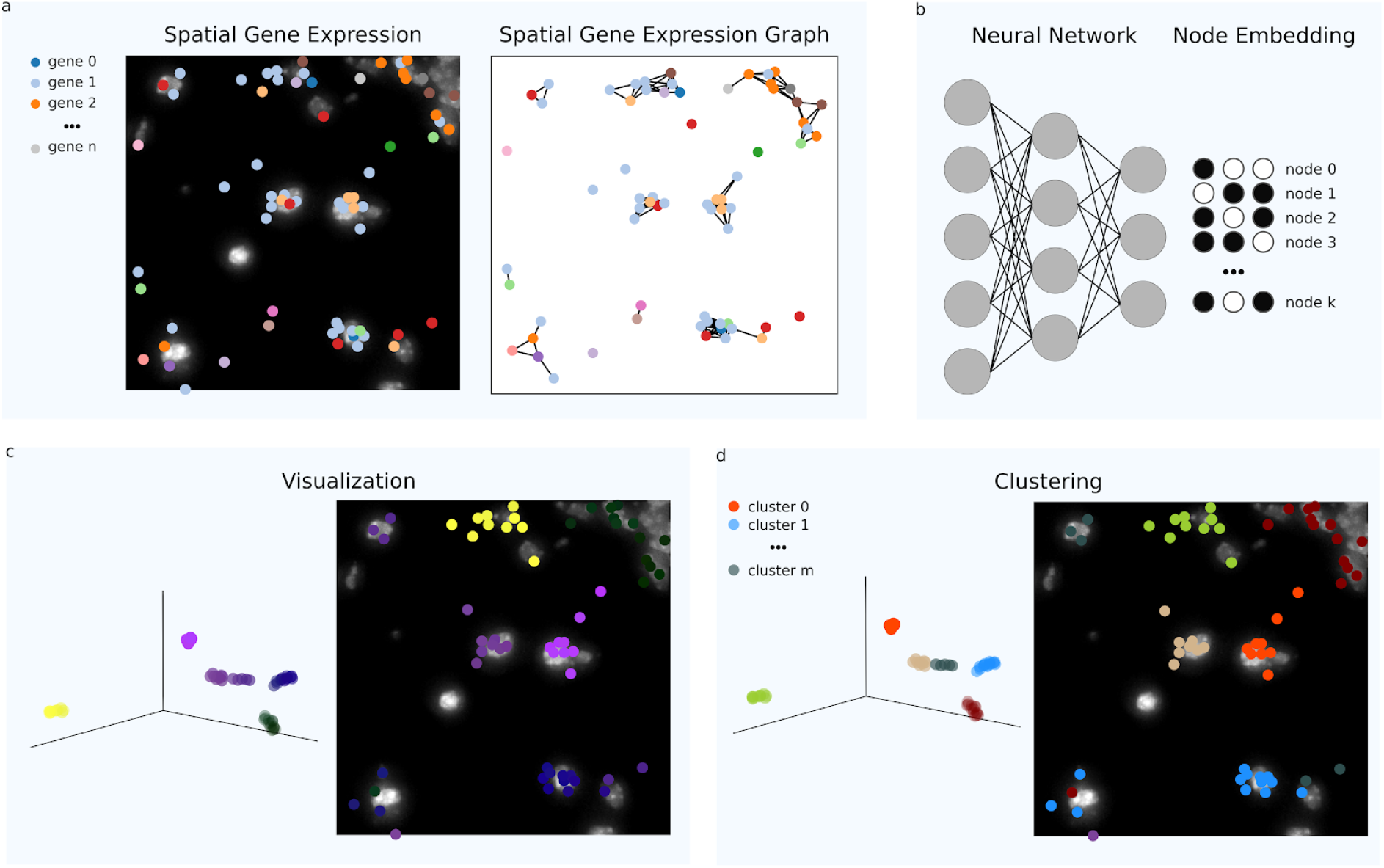
Spage2vec workflow for detecting subcellular spatial domains from spatial gene expression data. (**a**) Spatial transcript locations of *n* targeted genes are encoded in a graph connecting neighboring mRNA spots based on their spatial distances. (**b**) A lower dimensional representation is learnt for each of the *k* mRNA spots using a graph representation learning technique based on a graph neural network. The neural network predicts a node embedding vector for each mRNA of the graph representing high order spatial relationships with its local neighborhood (Materials & Method). Thereafter, the spatial gene expression variation can be (**c**) visualized at subcellular resolution projecting the learnt node embedding vectors in RGB color space, or (**d**) unsupervised clustering analysis can define *m* different clusters representing distinct subcellular spatial functional domains.

## RESULTS

### Spage2vec for in situ sequencing analysis

We first analyzed published in situ sequencing (ISS) data of mouse hippocampal area CA1 [11], where transcripts of 99 genes were localized. After representing the spatial gene expression as a graph, we applied spage2vec to generate a 50 dimensional embedding for each mRNA spot (Material & Methods), encoding information of its local neighborhood. We then projected the 50 dimensional embedding to three dimensions in order to visualize spatial relationships learnt from the data as similar colors in RGB color space [Figure 2a,c]. Next, in order to investigate if the learnt lower dimensional embedding contains significant information of biological functional domains, we clustered the spot embeddings directly in the 50-dimensional space (Material & Methods) and compared obtained spot cluster labels with cell-type annotations of spots from *Qian X. et al.* We initially obtained 29 clusters [Figure 2 supplementary 1], which reduced to 25 after merging highly correlated clusters (Material & Methods). Identified clusters can be interactively explored at https://tissuumaps.research.it.uu.se/demo/ISS_Qian_et_al.html [Supplementary File 1]. We then compared the 25 identified clusters with 20 cell-type- and 69 subcell-type-annotations defined in *Qian X. et al.*, excluding spots without cell-type labels [Figure 2e-f]. To demonstrate the ability of the model to generalize over unseen data, we used the spage2vec model trained on the right hemisphere mouse hippocampal area CA1 to predict the node embedding for the spatial gene expression graph of the left hemisphere CA1 area unseen during training [Figure 2b,d]. As can be seen in the figures [Figure 2a-d], the node representation of the two spatial gene expression graphs projected and visualized in RGB color space shows that the model produces visually similar embeddings for data not available during training.

**Figure 2.**
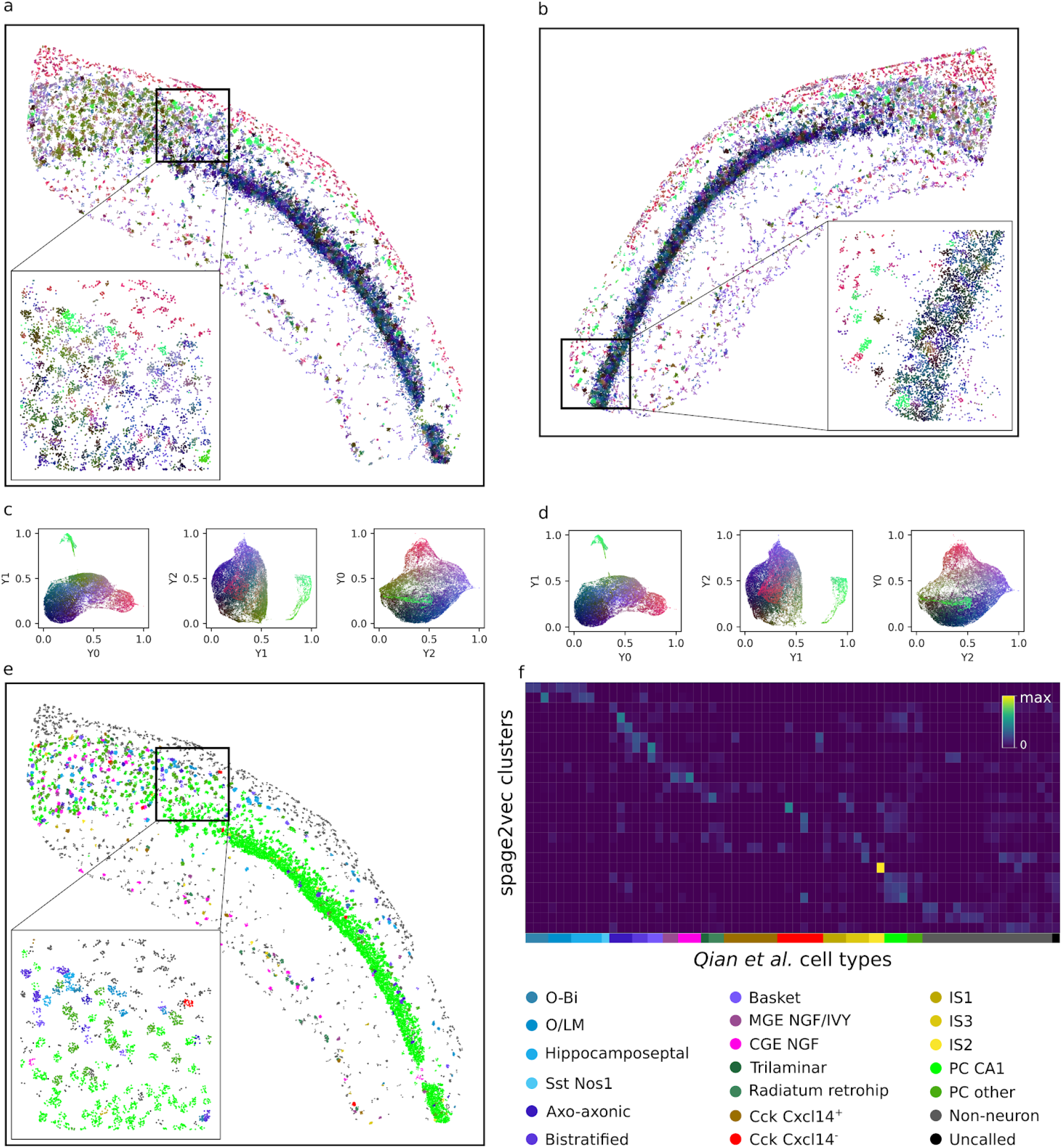
Application of spage2vec to in situ sequencing data of mouse hippocampal area CA1. Visualization of functional variation of spatial gene expression at subcellular resolution in right (**a**) and left (**b**) hippocampal area CA1 color coded based on their node embedding projections in RGB color space for right (**c**) and left (**d**) hemisphere. (**e**) Spatial gene expression with colored cell-type labels from *Qian X. et al.* analysis. (**f**) Heatmap showing the obtained spage2vec clusters with respect to cell- and subcell-type annotations (marked with different colors) from *Qian X. et al*., and cell-type legend.

### spage2vec for osmFISH analysis

In order to demonstrate the generalizability of spage2vec to other datasets, we also produced a lower dimensional representation of mRNAs from published osmFISH data of 33 cell-type marker genes targeted in mouse brain somatosensory cortex [7]. Again, we represented the gene expression as a graph and applied spage2vec, resulting in a 50 dimensional representation of each mRNA spot. We projected the 50 dimensions to three dimensions and visualized similar gene constellations as similar colors in 3D RGB color space [Figure 3a]. Next, we clustered the learnt embedding space in 274 domains [Figure 3 supplementary 1], and reduced to 69 domains after merging highly correlated clusters (Material & Methods). Identified clusters can be interactively explored at https://tissuumaps.research.it.uu.se/demo/osmFISH_Codeluppi_et_al.html [Supplementary File 1]. We then compared the resulting 69 clusters with the 31 cell-type annotations defined in *Codeluppi et al.*, excluding spots without cell-type labels [Figure 3b,c].

**Figure 3.**
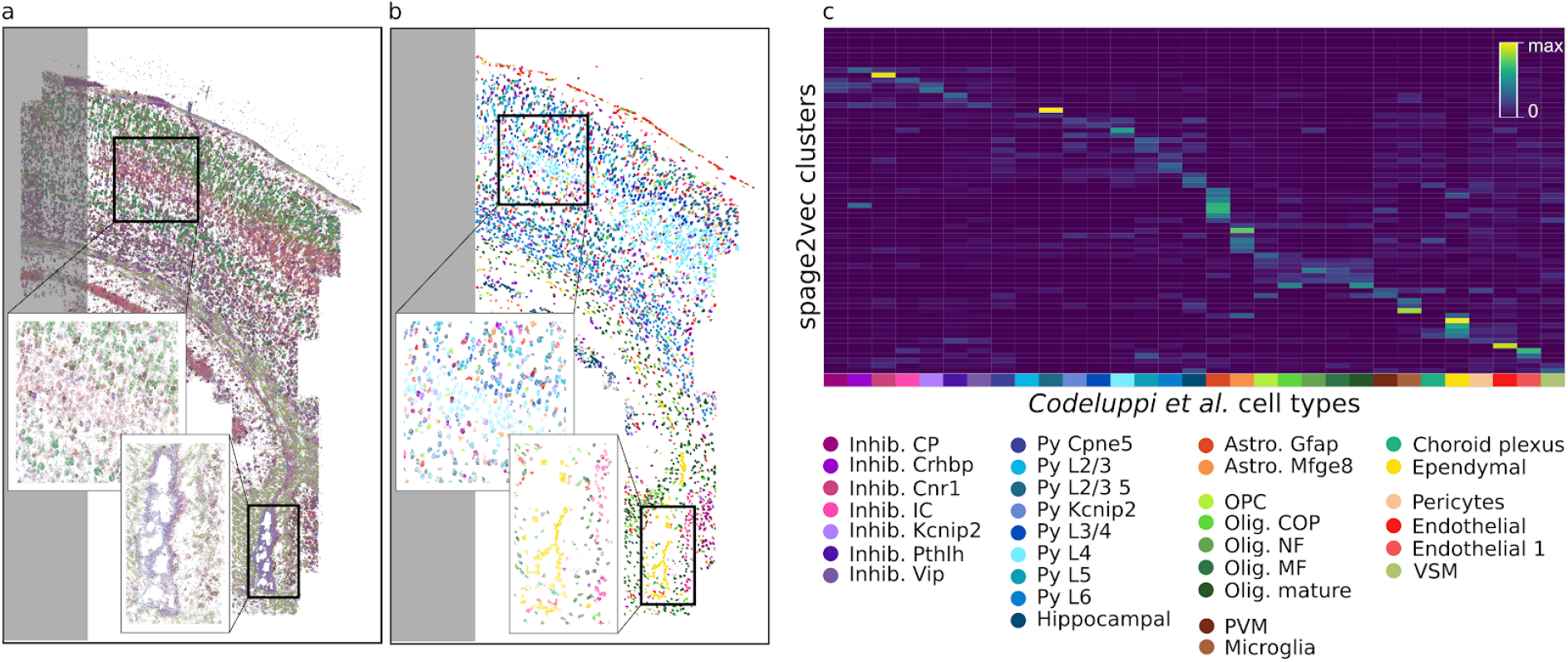
Application of spage2vec to osmFISH data from the mouse brain somatosensory cortex. (**a**) Visualization of functional variation of spatial gene expression at subcellular resolution color coded based on node embedding projection in RGB color space, and (**b**) spatial gene expression with colored cell-type labels from *Codeluppi S. et al.* cell segmentation. Shaded areas correspond to regions excluded in the original cell-type analysis. (**c**) Heatmap showing the obtained spage2vec clusters with respect to cell-type (marked with different colors) annotations from *Codeluppi S. et al*., and cell-type legend.

### Spage2vec for MERFISH analysis

We further applied spage2vec to a 3D mRNA localization dataset of hypothalamic preoptic region analyzed by MERFISH [6], where the transcripts of 135 targeted genes were localized in 3D. As for the previous dataset, we applied spage2vec to the graph representation (in this case 3D), and projected the 50 dimensions into three for visualization [Figure 4a]. Leveraging the symmetry of the data we trained a spage2vec model on approximately half the sample (0-956 µm) and tested on the other half. Clustering in 50-dimensional space resulted in 198 clusters [Figure 4 Supplementary 1], which reduced to 121 after merging of clusters with a gene expression correlation greater than 95%. Identified clusters can be interactively explored at *https://tissuumaps.research.it.uu.se/demo/MERFISH_Moffitt_et_al.html* [Supplementary File 1]. We compared the gene expression profiles of these 121 clusters with the 10 cell-types and 76 subcell-types presented in [6] [Figure 4b-d].

**Figure 4.**
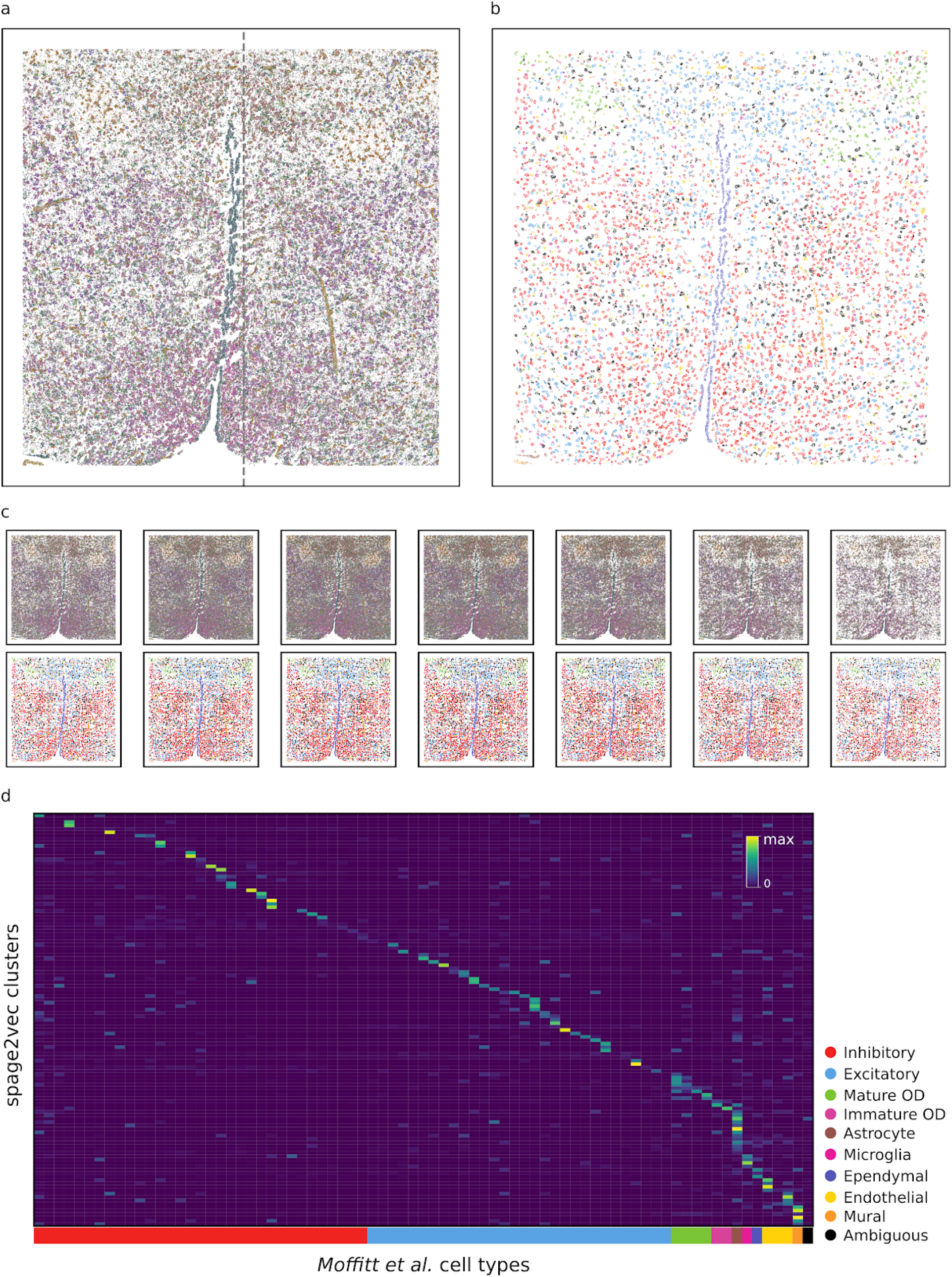
Application of spage2vec to MERFISH data of the mouse brain hypothalamic preoptic region. (**a**) Visualization of functional variation of spatial gene expression at subcellular resolution color coded based on their node embedding projections in RGB color space. The gray dashed line defines regions of the sample used for training (left) and for testing (right). (**b**) Spatial gene expression with colored cell-type labels from *Moffitt J. R. et al.* cell segmentation. (**c**) Spatial distribution of node embedding projections in RGB color space (upper row) and cell-type labels (bottom row) from *Moffitt J. R. et al.* across the whole section. (**f**) Heatmap showing the obtained spage2vec clusters with respect to cell- and subcell-type annotations (marked with different colors) from *Moffitt J. R. et al*., and cell-type legend.

## DISCUSSION

We showed that spage2vec can learn low dimensional embeddings encoding important topological and functional information of local gene expression.This rich low dimensional space can be used for downstream clustering analysis in order to detect biologically meaningful re-occuring gene constellations that correlate well with subcellular and cellular domains. The embedding, found by unsupervised training, has an inductive property to generalize over unseen nodes. This means that it can be applied to a new unseen dataset, as long as the new dataset has the same feature set (e.i., consists of gene expression data from the same gene panel). This is especially useful to predict embeddings for new spatial gene expression datasets and map them to a common lower dimensional space. The fact that spage2vec is a fully unsupervised approach triggers the possibility for the discovery of novel cell-types in situ without the need of scRNA sequencing data driven analysis.

The presented approach is completely independent of cell segmentation, and equally applicable to 2D and 3D data, meaning that dense gene expression datasets such as those from MEHRFISH can be analyzed without relying on the accuracy of cell segmentation. In fact, most cell segmentation approaches are based on identifying cell nuclei, and then approximating gene-to-cell assignment by shortest distance to the closest nucleus. This can very often introduce noise as cells may vary very much in shape, and the nucleus of a given cell may not even be present in the same tissue section as the bulk of the cell. Furthermore, the presented segmentation free spage2vec approach enables detection of cell types with varying sub-cellular gene expression patterns as well as subcellular constellations of genes representing functional domains located far away from a cell nucleus.

## MATERIAL & METHODS

### Building a Spatial Gene Expression Graph

Spatially resolved gene expression data consists of gene expression information and coordinates describing spatial location (in 2D or 3D) in a tissue sample. This information can be represented as a graph by saying that a node in the graph has a single categorical feature representing the gene expression (mRNA) it belongs to. Next, connections are drawn between each node and all its local neighbors within a maximum spatial distance *d*_*max*_. The distance *d*_*max*_ is defined such that at least 97 percent of all nodes are connected to at least one nearest neighbor, automatically adjusting for the spatial resolution of the dataset. Connected components with less than three nodes representing spurious expressions are removed from the graph before further processing [Figure 1a]. Note that the same graph representation works in both 2D and 3D.

### Neural Network Model and Training

Next, spage2vec strives to transform the spatial gene expression graph into an embedding where similar gene constellations are assigned similar vectors using a neural network model. The neural network model consists of an unsupervised GraphSAGE [27] model implemented with the open source machine learning python library StellarGraph [29]. The model learns embeddings of unlabeled graph nodes by combining the node’s own feature with features sampled and aggregated from the node’s local neighborhood. Specifically, node embeddings are learnt by solving a binary node classification task that predicts whether arbitrary node pairs are likely to co-occur in a random walk performed on the graph. For this task the training set consists of *positive* node pairs, pairs that co-occur within walks of length 2 on the graph, and *negative* pairs of nodes uniformly randomly selected from the graph. Through training this binary node pair classifier, the model automatically learns an inductive mapping from a high-dimensional feature space (i.e. spatial gene expression) to a lower dimensional node embedding space, describing gene constellations, preserving important topological and structural features of the nodes. The model architecture consists of two identical GraphSAGE encoder networks sharing weights, taking as input a pair of nodes together with the graph structure and producing as output a pair of node embeddings. Thereafter, a binary classification layer with a sigmoid activation function, learns to predict how likely it is that a pair will occur at a random position in the graph. Model parameters are optimized by minimizing binary cross-entropy between the predicted node pair labels and the true labels, without supervision.

### Neural network hyperparameters

The proposed spage2vec model architecture used for all experiments presented here consists of two GraphSAGE layers with 50 hidden units, a bias term, l2 normalization, and l1 kernel regularization, using attentional aggregator function [30] with LeakyRelu [31]. Each GraphSAGE encoder embeds each node’s neighborhood with a 2-hop node aggregation strategy, sampling respectively 20 and 10 nodes for the first and the second hop. The model is trained with on-the-fly batch generation with batch size equal to 50, using Adam [32] as optimizer with learning rate equal to 0.5e-4. The output of spage2vec will thus be one vector of length 50 per spatial gene expression position. All details and settings are provided as Python notebooks (https://github.com/wahlby-lab/spage2vec).

### Visualization of node embeddings

To visualize the extracted spatial gene expression embeddings created by spage2vec, we reduced the embedding dimensionality to three dimensions with UMAP [33]. This allowed us to present the spatial gene expression constellations as data points in a 3D RGB color space. Mapping the new color-coding back to tissue space shows that many of the constellations not only cluster in space but also seem to recur and correlate with cellular and subcellular spatial domains [Figure 1d].

### Identification of distinct gene constellations and spatial domains

For further comparing the spage2vec output with approaches aimed at identifying cell types we hypothesize that recurring constellations of genes are spatial functional domains that may be cell type specific, or represent processes shared among different cell types. We therefore cluster the 50-dimensional spage2vec output using the Leiden clustering algorithm [34,35] followed by Z-score normalization of the cluster expression matrix (cluster x genes). Clusters where gene expression counts have a correlation greater than 95% are merged, and the merged cluster expression matrix is re-normalized with Z-score normalization, leading to a final set of clusters. Note that the trained model has an inductive property, meaning that it can generalize and find embeddings for previously unseen gene constellations.

### Datasets

We apply spage2vec to three publicly available published mouse brain tissue datasets obtained by three different spatial transcriptomics assays: (1) In situ sequencing (ISS) of left and right hippocampal area CA1 [11, https://tissuumaps.research.it.uu.se/demo/ISS_Qian_et_al.html], with a resolution of 0.325 μm per px and a total of 84880 detections of 99 different mRNAs. We refer to this as the ISS dataset. (2) An osmFISH analysis of the somatosensory cortex [7, https://tissuumaps.research.it.uu.se/demo/osmFISH_Codeluppi_et_al.html], comprising a tissue section of 3.8 mm^2^, with a resolution of 0.065 μm per pixel, and a total of 1802589 detections of 33 different mRNAs. We refer to this as the osmFISH dataset. (3) A MERFISH analysis of the hypothalamic preoptic region [6, https://tissuumaps.research.it.uu.se/demo/MERFISH_Moffitt_et_al.html], comprising a 3D tissue section 10 μm thick of 1.8 by 1.8 mm and a total of 3728169 detections targeting 135 different genes, referred to as the MERFISH dataset.

### Code Availability

All software was developed in Python 3 using open source libraries. The processing pipeline and the source code used to generate figures and analysis results presented in this paper are available as Python notebooks at https://github.com/wahlby-lab/spage2vec.

## ACKNOWLEDGMENTS

We thank Mats Nilsson, Sten Linnarsson and Xiaowei Zhuang for making their datasets publicly available. We also thank Leslie Solorzano for providing support in visualization of the results with the TissUUmaps viewer. This research was funded by the European Research Council via ERC Consolidator grant 682810 to C. Wählby and Swedish Foundation for Strategic Research (grant BD150008).

## COMPETING INTERESTS

The authors have no competing interests.

---

**Figure 2 supplement 1.**
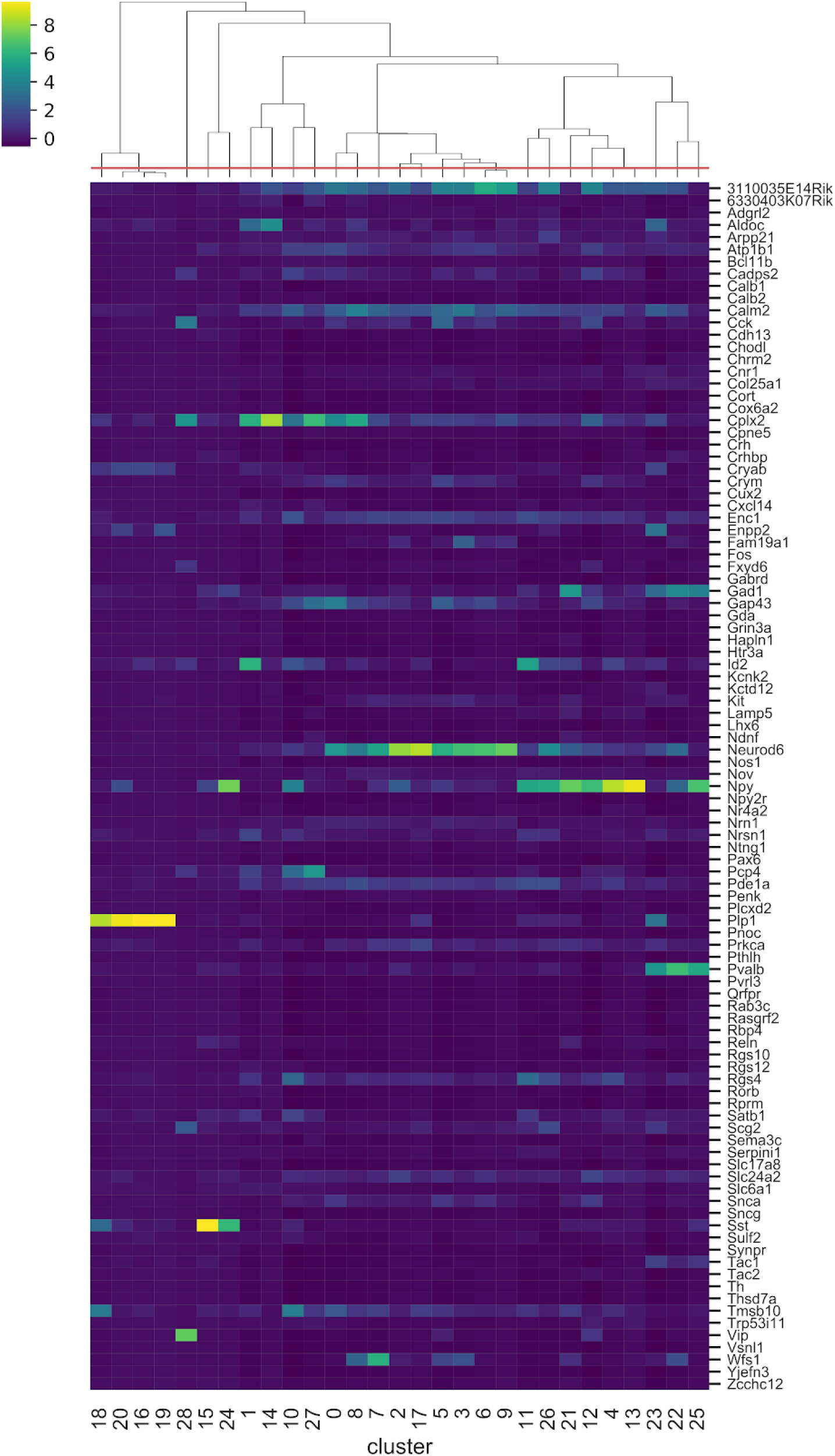
Gene expression per detected cluster, or gene constellation. Each column represents a cluster from the spage2vec embedding of the ISS data from *Qian X. et al.* and each row shows how much each gene contributes to a given cluster with Z-score normalized values. The red line on top of the dendrogram shows the correlation threshold used for merging clusters.

**Figure 3 supplement 1.**
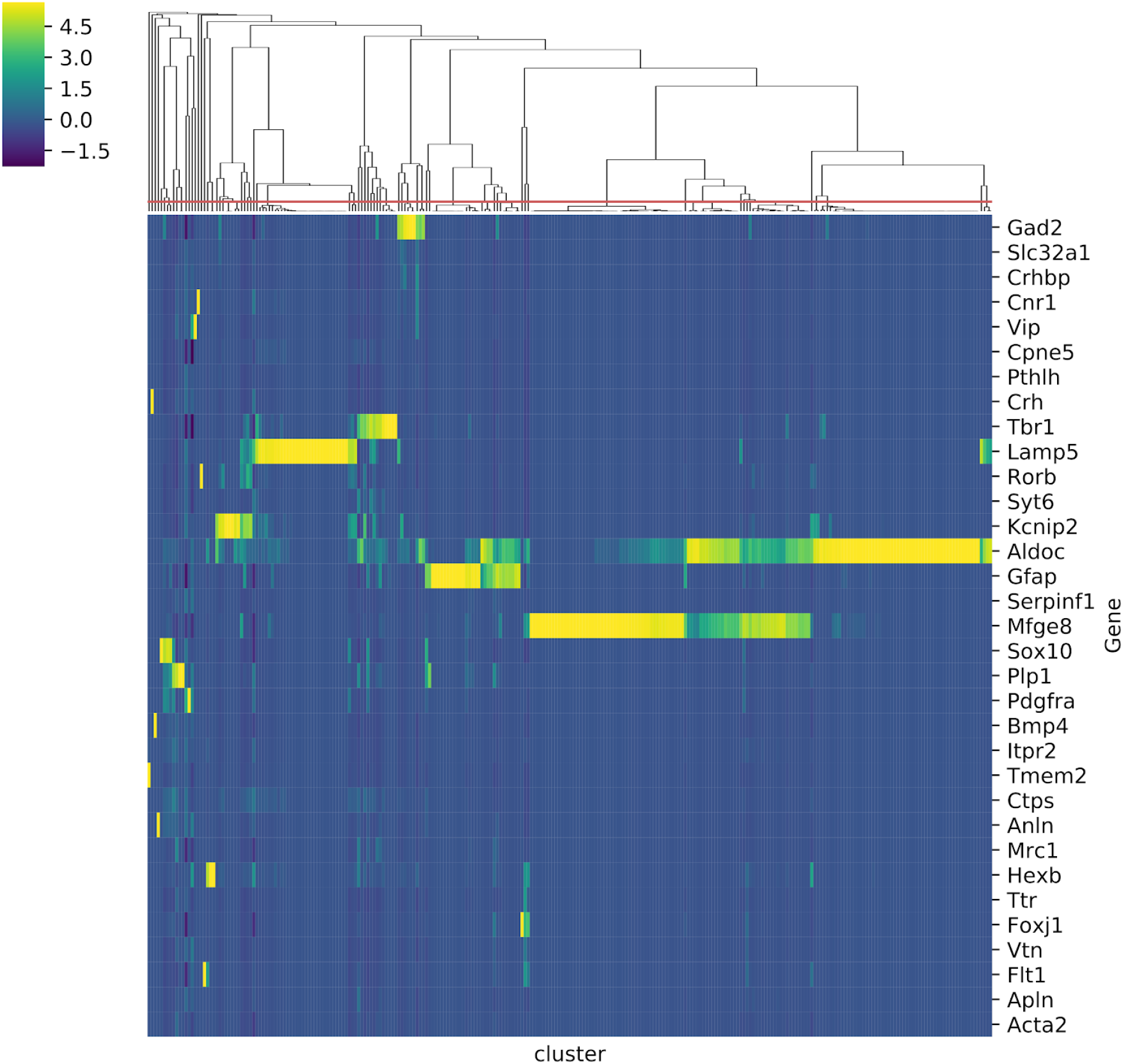
Gene expression per detected cluster, or gene constellation. Each column represents a cluster from the spage2vec embedding of the osmFISH data from *Codeluppi S. et al*., and each row shows how much each gene contributes to a given cluster with Z-score normalized values. The red line on top of the dendrogram shows the correlation threshold used for merging clusters.

**Figure 3 supplement 1.**
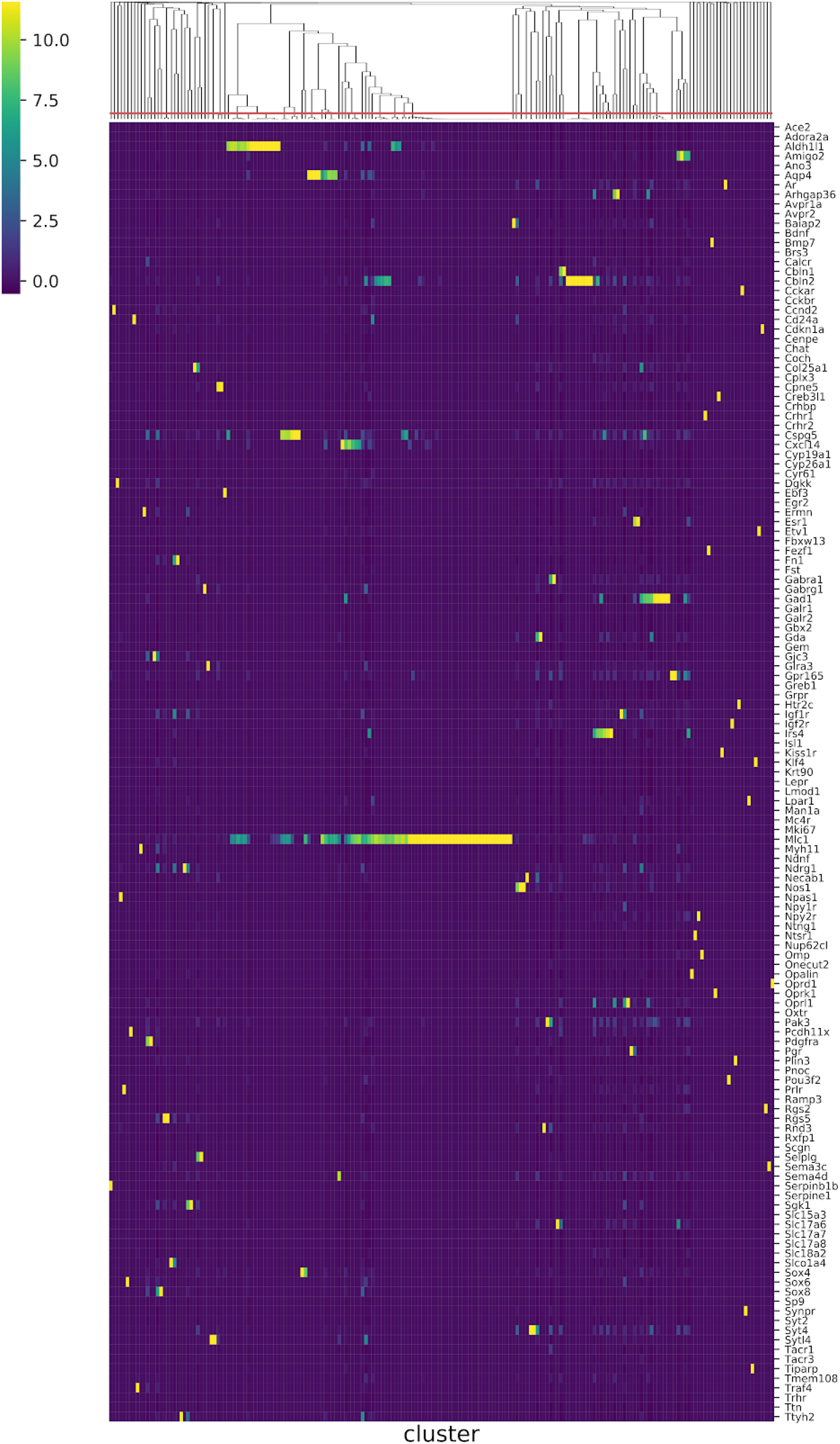
Gene expression per detected cluster, or gene constellation. Each column represents a cluster from the spage2vec embedding of the MERFISH data from *Moffitt J.R. et al*., and each row shows how much each gene contributes to a given cluster with Z-score normalized values. The red line on top of the dendrogram shows the correlation threshold used for merging clusters.

## SUPPLEMENTARY FILE 1

### Visualization of spage2vec clusters in TissUUmaps online viewer

1. Open in a browser one of the following websites:
  - ISS dataset: *https://tissuumaps.research.it.uu.se/demo/ISS_Qian_et_al.html*
  - osmFISH dataset: *https://tissuumaps.research.it.uu.se/demo/osmFISH_Codeluppi_et_al.html*
  - MERFISH dataset: *https://tissuumaps.research.it.uu.se/demo/MERFISH_Moffitt_et_al.html*
2. Click on `Download data` in `Marker data` -> `Gene expression` tab, analysis results will load in your browser.
3. Select “`macro_cluster”` from `cluster column` drop down menu
4. Select “`global_X_pos`” from `X column` drop down menu
5. Select “`global_Y_pos`” from `Y column` drop down menu
6. Click on `Load markers`, the list of clusters with read counts, color and marker shape will appear.
7. Check the `Show` box of the clusters you wish to visualize *Note: For efficient visualization at the lower magnifications only a fraction of reads will be displayed, while the number of displayed markers will increase zooming in to the highest magnification (displaying all markers in the field of view)*.
8. Marker size can be changed in `Global size` box for all the markers or in the `size` box for the individual marker, as well as marker color and shape. Zooming in or out will refresh the view and the update will be in place.

## Notes

https://tissuumaps.research.it.uu.se/spage2vec/index.html

